# Spatio-temporal monitoring of lake fish spawning activity using environmental DNA metabarcoding

**DOI:** 10.1101/2022.02.07.478003

**Authors:** C. Di Muri, L. Lawson Handley, C. W. Bean, M. Benucci, L. R. Harper, B. James, J. Li, I. J. Winfield, B. Hänfling

## Abstract

Determining the timing and location of fish reproductive events is crucial for the implementation of correct management and conservation schemes. Conventional methods used to monitor these events are often unable to assess the spawning activity directly or can be invasive and therefore problematic. This is especially the case when threatened fish populations are the study subject, such as the Arctic charr (*Salvelinus alpinus* L.) populations in Windermere (Cumbria, UK). Arctic charr populations have been studied in this lake since the 1940s, and the locations and characteristics of spawning grounds have been described in detail using techniques such as hydroacoustics, as well as physical and visual surveys of the lake bottom. Here, in conjunction with established netting surveys, we added an environmental DNA (eDNA) metabarcoding approach to assess the spatial distribution of Arctic charr in the lake throughout the year to test whether this tool could allow us to identify spawning locations and activity. Sampling was carried out between October 2017 and July 2018 at three locations in the lake, covering putative and known spawning sites. eDNA metabarcoding provided accurate spatial and temporal characterisation of Arctic charr spawning events. Peaks of Arctic charr read counts from eDNA metabarcoding were observed during the spawning season and at specific locations of both putative and known spawning sites. Net catches of mature Arctic charr individuals confirmed the association between the Arctic charr spawning activity and the peaks of eDNA metabarcoding read counts. This study demonstrates the ability of eDNA metabarcoding to effectively and efficiently characterize the spatial and temporal nature of fish spawning in lentic systems.

## Introduction

Anthropogenic pressures are threatening freshwater fish populations worldwide (Díaz *et al*., 2019), and conservation biologists and environmental managers are striving to preserve such diversity as it provides ecosystem services to humans and holds intrinsic evolutionary and ecological value (Lynch *et al*., 2016; Piccolo, 2017). Arctic charr (*Salvelinus alpinus* L.) has the most northerly distribution of all anadromous freshwater teleosts (Hansen *et al*., 2019), ranging from the temperate areas of eastern North America and the European Alps, to the most northern points of the Eurasian and North American continents (Johnson, 1980). The species is adapted to cold, highly oxygenated waters and is therefore especially vulnerable to climate change and eutrophication (Winfield *et al*., 2008). As a consequence, many local populations of Arctic charr have already become extinct (Kelly *et al*., 2020; Winfield *et al*., 2010). Despite the observed extinctions and population declines, the level of national and international protection is low. Globally, Arctic charr are not considered to be threatened (Freyhof and Kottelat, 2008), but within Great Britain and Ireland, where the species are at the western edge of their European range, their presence is considered to be rare. Whilst this species does not appear in the annexes of the European Union Habitats and Species Directive (Adams *et al*., 2007), Arctic charr has been included as a priority conservation taxon within the UK Biodiversity Action Plan and a high conservation value species because of its limited distribution, past extirpations and current concerns over the conservation status of many populations (Maitland *et al*., 2007; Bean *et al*., 2018).

Interest in Arctic charr conservation is primarily driven by the often-unique characteristics of individual populations. At the onset of the most recent post-glacial era, the isolation of many Arctic charr populations in post-glacial lakes led to high polymorphism and plasticity among and within local populations (Jonsson and Jonsson, 2001; Klemetsen, 2010; Skúlason*et al*., 2019). This high diversity reflects the ability of Arctic charr populations to adapt to cold, nutrient impoverished and species-poor lacustrine habitats with a variety of niches available (Jonsson and Jonsson, 2001; Reist *et al*., 2013). Arctic charr polymorphism encompasses life-history tactics (e.g. anadromous and non-migratory forms; Klemetsen *et al*., 2003), specialisation in diet and habitat preferences associated to morphological variation (Adams *et al*., 2003; Klemetsen *et al*., 2006), and reproductive strategies (Frost, 1965; Smalås, Amundsen and Knudsen, 2013; Telnes and Saegrov, 2004). For example, in lakes where sympatric populations of Arctic charr exist, differences in the spatial and temporal separation of spawning grounds can occur, leading to the emergence of phenotypic differences and genetic divergence between populations (Garduño-Paz *et al*., 2012).

Arctic charr typically spawn for a limited period at shallow gravel banks where the females dig depressions in which eggs are incubated (Esteve, 2005). Here, a clean substrate with low amounts of fine sediment and a well-oxygenated interstitial zone, is required to ensure successful reproduction (Sternecker, Denic and Geist, 2014). These stringent requirements mean that only a small fraction of available lake habitat is typically used for spawning. Low *et al*. (2011) estimated that in Irish lakes, Arctic charr spawning substrate comprised between just 0.4-0.7% of the littoral habitat, and that egg numbers are significantly correlated with gradient and spawning site width. Furthermore, a number of studies have recognized that siltation and sedimentation on the spawning gravels are major causes of reproductive failure (Franssen *et al*., 2012; Levasseur *et al*., 2006; Winfield *et al*., 2010).These variables, but often very specific, temporal and spatial characteristics of Arctic charr spawning mean that detailed local knowledge is invaluable for conservation efforts, to ensure protection of high-quality habitats for reproduction and to identify management units.

Arctic charr spawning areas and spawning behavior in Windermere (Cumbria, UK) have been extensively studied over more than 50 years (Frost, 1965; Miller *et al*., 2015). Research on breeding habitats of Windermere’s Arctic charr has described the presence of two sympatric populations with autumn and spring spawning events (Frost, 1965), and documented their patterns of genetic and phenotypic divergence (Corrigan *et al*., 2011; Partington and Mills, 1988). Outside the spawning season, Arctic charr in Windermere are exclusively restricted to offshore areas of the lake (Lawson Handley *et al*., 2019; Winfield *et al*., 2008). Autumn-spawners release their gametes at depths of around 2 m between November and December, whereas spring-spawners mature between February and March and spawn at deeper sites between 15 and 20 m. Both the mesotrophic north basin and eutrophic south basins of Windermere sustain autumn and spring-spawners, and a variety of putative and known spawning locations have been described in the lake (Frost 1965; Miller *et al*., 2015; Winfield *et al*., 2015). However, these Windermere Arctic charr populations have declined markedly over the last few decades (Winfield *et al*., 2019) in parallel with increased eutrophication and the associated decrease in spawning habitat quality (Miller *et al*., 2015; Winfield *et al*., 2015). Suitable spawning habitats are now limited to the shallowest areas of the lake at depths below 5 m where clean, hard substrates still occur (Miller *et al*., 2015, Winfield *et al*., 2015). This poses important conservation concerns for spring-spawning Arctic charr populations with deeper breeding grounds (Winfield *et al*., 2008). Importantly, extensive monitoring of spawning activity at depths less than 5 m is challenging using established non-invasive survey methods such as hydroacoustic applications (Miller *et al*., 2015), and netting surveys cannot be deployed widely due to the conservation status of target species. Non-invasive and broadly applicable monitoring approaches are therefore required to characterise times and locations of fish spawning activities.

The analysis of environmental DNA (eDNA) has been recently applied to the study of riverine fish spawning activity whereby the isolation of genetic material from water samples, eDNA, coupled with species-specific quantitative PCR assays, allowed spawning aggregation and migration patterns of riverine fish to be identified (e.g. Antognazza *et al*., 2019; Bracken *et al*., 2019; Thalinger *et al*., 2019). Temporally and spatially constrained changes in eDNA concentration of target fish species were found to be associated with spawning activity and release of gametes. However, this approach has not yet been applied to fish populations in a standing water.

Recent studies have demonstrated that eDNA metabarcoding, which is used to describe the entire fish community as opposed to single target species, is able to detect seasonal variation in community composition, heterogeneity in the use of habitat, and even estimates of population biomass and abundance (Di Muri *et al*., 2020; Lawson Handley *et al*., 2019). Here we apply eDNA metabarcoding to investigate the spawning activity of Arctic charr in Windermere. Based on evidence provided by previous eDNA research, we hypothesise that (1) eDNA metabarcoding analyses, using extensive spatio-temporal water sampling, can detect lake fish spawning activity via the temporal and spatial variation in eDNA read counts during the breeding season, and that (2) species-specific peaks in read counts from eDNA metabarcoding reflect the sites and times where spawning events are expected. To test our hypotheses, we focused on the shallowest breeding grounds of the autumn-spawning Arctic charr population in Windermere’s north basin that have recently been assessed as being suitable to support spawning activity (Miller *et al*., 2015; Winfield *et al*., 2015). Specifically, we targeted putative and known spawning grounds, with the latter monitored by annual netting since 1940 (Winfield *et al*., 2008). Gill-netting survey results were compared with Arctic charr read counts associated with spawning individuals caught at these sites. Additionally, in autumn, we expected an eDNA signal of similar strength at the putative spawning grounds if any spawning activity was occurring.

## Materials and methods

### Water sample collection, filtration, and DNA extraction

Water sampling was carried out in Windermere (UK, Fig. 1) on twelve dates between October 2017 and July 2018, with a higher sampling effort in late autumn when Arctic charr are expected to be reproductively active (Fig. 2). Water samples were collected at the known autumn spawning site of North Thompson Holme island (NTH, three locations), and along two transects located at the putative spawning site on the west shore at Red Nab (RN, eight locations) and at offshore locations approximately in the middle of the lake (OF, five locations) that are deep-water feeding habitats (Fig. 1). Due to logistic reasons (i.e. suitable boat not available) we were unable to collect water samples at NTH on the 01^st^, 14^th^ and 16^th^ of November, and RN water samples were not collected on the 08^th^ of November (Fig. 2). Site coordinates were recorded during the first eDNA sampling event (18^th^ October 2017) using a hand-held Geographic Positioning System (GPS) (Garmin eTrex 10, Kansas, USA; Table S1) and these coordinates were used to navigate to the collection sites during all subsequent events. At NTH between October and December 2017, spawning activity at the time of sampling was verified by catches of Arctic charr individuals from twelve gill-netting surveys. The maturity of Arctic charr individuals caught at NTH was assessed via the morphological evaluation of body shape and coloration as well as the ease with which gametes could be expressed. At RN, Arctic charr spawning activity has not been recently monitored or demonstrated through catches of spawning individuals. However, the area has been identified as a putative spawning ground based on anecdotal historical records, presence of suitable substrate, and passive acoustic monitoring (PAM) data which identified noises connected to Arctic charr spawning activities (gravel displacement or sounds associated with air exchange with swimbladder regulation; Bolgan *et al*., 2018).

**Figure 1.**
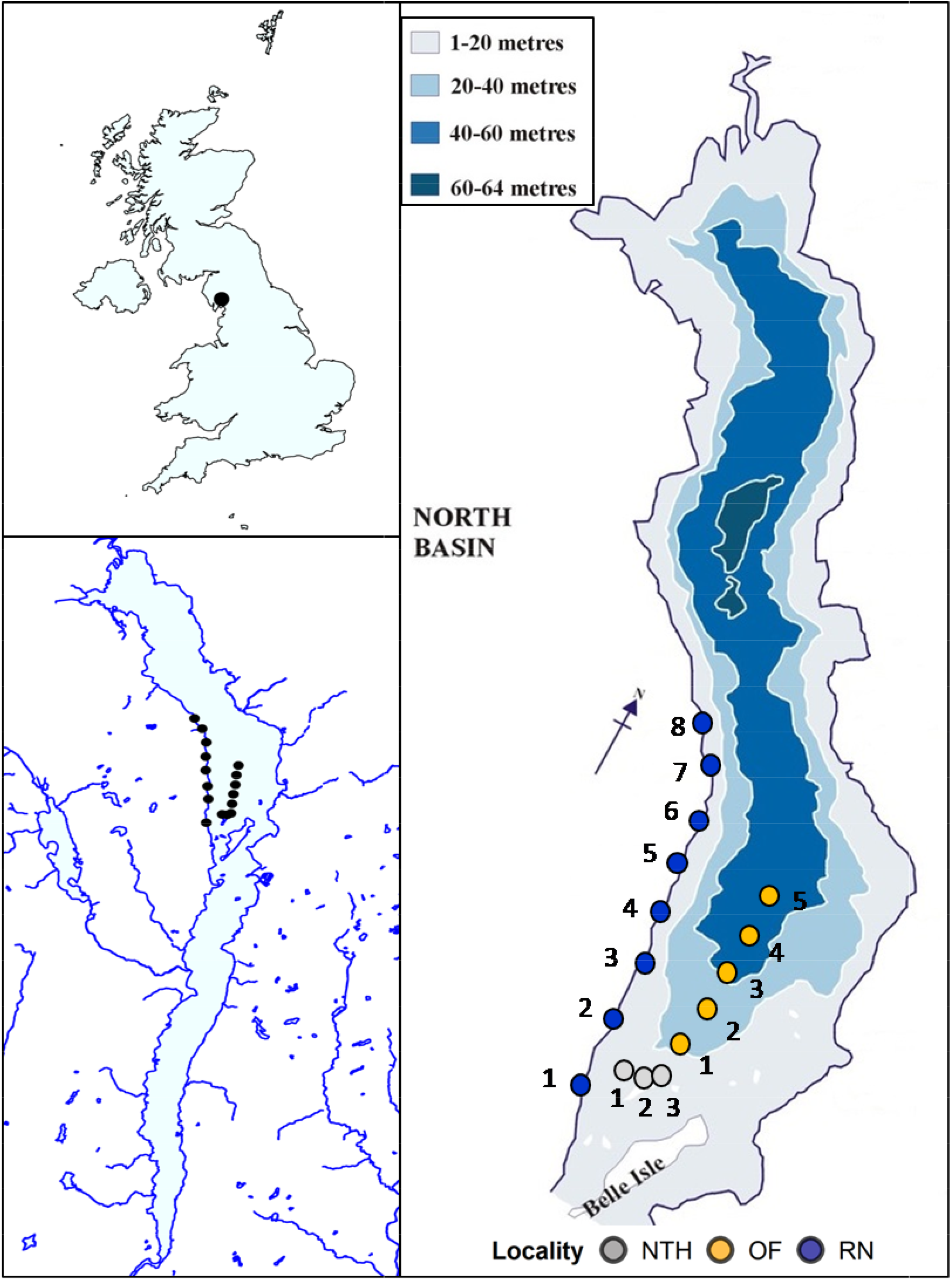
Map of Windermere and eDNA collection sites. Windermere’s location in Cumbria, UK and location of eDNA sampling sites in Windermere’s north basin (left side panels). Detailed bathymetric map of Windermere’s north basin with sites and localities sampled during our eDNA surveys (right side panel). In the bathymetric map, “OF” are offshore sites (Arctic charr feeding grounds), “NTH” are inshore sites located at the shore of North Thompson Holme island (Arctic charr monitored spawning grounds) and “RN” are shoreline sites on the west side of the lake at Red Nab (Arctic charr putative spawning grounds). The bathymetric map was edited from Ramsbottom (1976) and used with permission of the Freshwater Biological Association.

**Figure 2.**
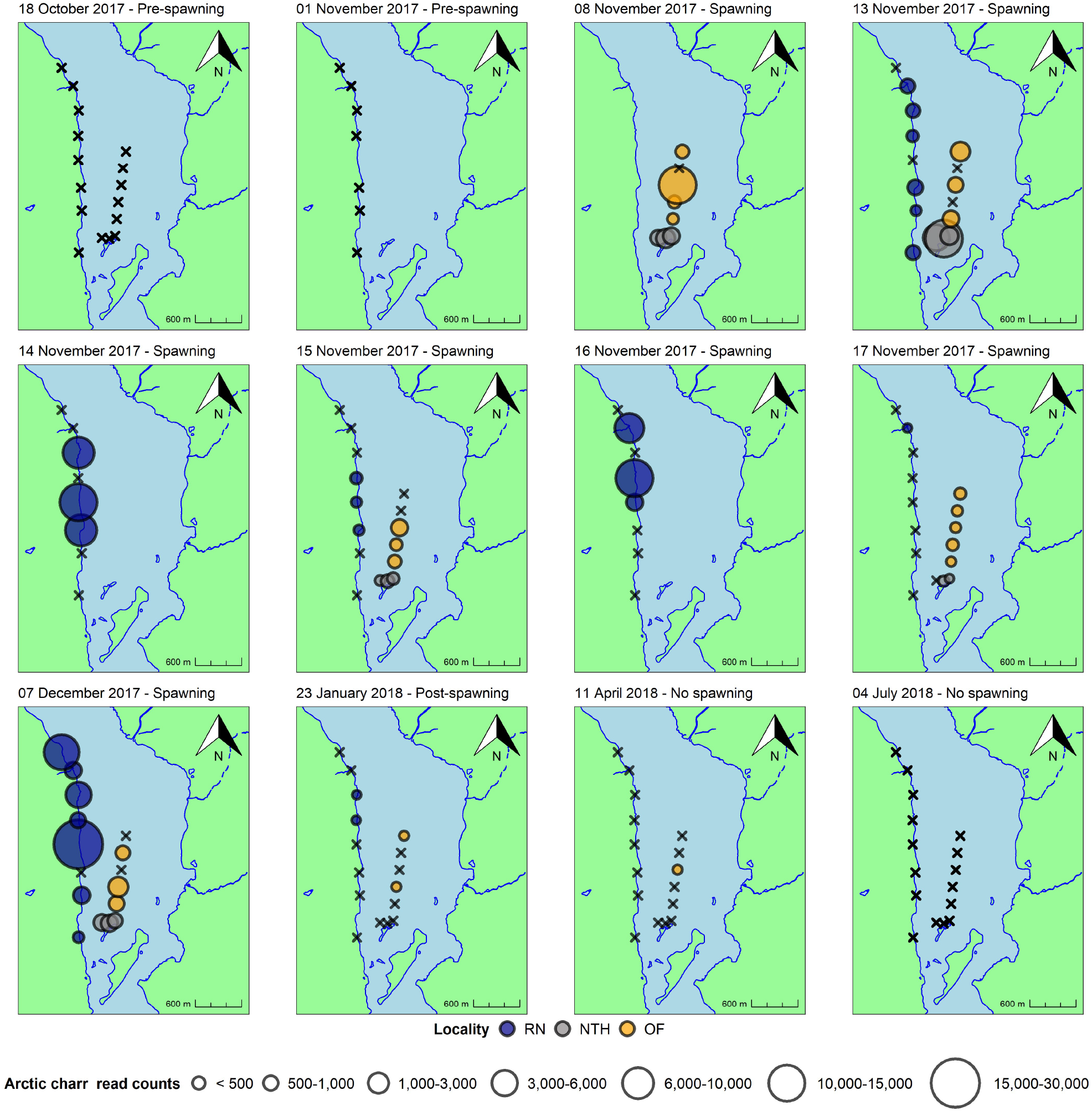
Spatio-temporal variation of Arctic charr eDNA signal in the north basin of Windermere. Bubble size is proportional to read counts assigned to Arctic charr, whereas black crosses indicate sites where samples were collected but the species was not detected, and the absence of any symbol indicates that water samples were not collected.

At each of the eight sites in the onshore location (RN; less than 0.5 m depth), five subsamples (5 × 400 mL taken across 50 m) were collected at the surface water layer from the shoreline and merged into a single 2 L sterile plastic bottle (Gosselin™ Square HDPE, Fisher Scientific UK Ltd, UK). One sampling blank, consisting of a 2 L sterile bottle filled with ultra-purified water (Milli-Q), was used and opened once in the field. In the middle-lake sites, NTH and OF, samples were collected at different depths (2 m to 40 m; Fig. 1) from a boat using a 1.5 L, Friedinger-like water sampler left semi-open to fill up during the descent. The water sampler was lowered three times at each sampling site (3 x ∼650 mL) in order to collect subsamples that were subsequently merged into a single 2 L sterile plastic bottle. The water sampler was sterilised, while moving between sites, by soaking in a 10% v/v chlorine-based commercial bleach solution (Elliott Hygiene Ltd, UK) for 10 minutes, followed by rinsing with 5% v/v MicroSol detergent (Anachem, UK) and purified water. At each site, after sterilisation, the sampler was also quickly lowered and washed with the lake’s water before collection occurred. Two sampling blanks were used during the offshore sample collection (beginning and end of water sampling) to account for potential contamination introduced by the use of a water sampler. After bleaching, the Milli-Q water of each blank was used to rinse the water sampler before pouring it back into the 2 L bottles. To further minimise contamination risk, nitrile sterile gloves (STARLAB, UK) were worn and changed between collection sites.

Water samples were kept in cool boxes covered with ice packs and filtered within six hours after water collection. Water was filtered using vacuum-pumps coupled with Nalgene™ units and DNA was captured onto 0.45 μm mixed cellulose ester filters (47 mm diameter, Whatman, GE Healthcare). Generally, two filters were used per 2 L of water collected from the shoreline sites and one filter was used for water samples collected offshore with a few exceptions. A filtration blank was run during each filtration round (*n* = 21), where 2 L of ultra-purified water was filtered alongside water samples (*n* = 160) and sampling blanks (*n* = 29). Filters were stored in sterile 50 mm Petri dishes (Fisher Scientific UK Ltd, UK), sealed with parafilm (Bemis™, Fisher Scientific UK Ltd, UK), at -20° C until extraction.

The mu-DNA water protocol (Sellers *et al*. 2018) was used for DNA extraction from filters, and filters from the same sample were lysed together in the same tube. Samples, sampling blanks, and filtration blanks belonging to different sampling dates were extracted in separate batches. An extraction blank, consisting only of extraction reagents, was included for each extraction round (*n*=13). DNA yield and purity were checked using a Nanodrop 1000 spectrophotometer (Thermo Fisher Scientific).

### Sequencing library preparation

The sequencing library was built using a custom library preparation protocol which includes tagged primers in two rounds of PCR (*sensu* Li *et al*., 2019). For the first PCR, indexed primers amplifying a ∼106 bp region of the mitochondrial 12S ribosomal RNA (rRNA) in fish were used (Kelly *et al*. 2014; Riaz et al. 2011). PCR was performed with a final reaction volume of 25 μL, including 12.5 μL of Q5^®^ Hot-Start High-Fidelity 2X Master Mix (New England Biolabs^®^ Inc., MA, USA), 1.5 µL of each indexed primer (10 µM; Integrated DNA Technologies, Belgium), 0.5 µL of the Thermo Scientific™ Bovine Serum Albumin (Fisher Scientific UK Ltd, UK), 7 µL of molecular grade water (Fisher Scientific UK Ltd, UK) and 2 μL of DNA template at the original sample concentration. To avoid cross-contamination between samples, reactions were prepared in 8-strip tubes with individually attached caps and covered with a drop of mineral oil (Sigma-Aldrich Company Ltd, UK). Amplifications were performed on Applied Biosystems^®^ Veriti thermal cyclers (Life Technologies, CA, USA) with the following conditions: initial denaturation at 98°C for 5 min; 35 cycles of 98°C for 10 sec, 58°C for 20 sec and 72°C for 15 sec; final elongation at 72°C for 7 min. Samples, blanks, PCR negative controls (molecular grade water, *n* = 11), and PCR positive controls (genomic DNA [0.05 ng/µL] from *Maylandia zebra*, a cichlid from Lake Malawi not present in UK, *n* = 11) were amplified in triplicate. Amplicons were checked on 2% agarose gels stained with 10,000X GelRed Nucleic Acid Gel Stain (Cambridge Bioscience, UK). Gels were imaged using Image Lab Software (Bio-Rad Laboratories Ltd, UK) to visually check for contamination in blanks/PCR negative controls, presence of target band, and consistency of results among replicates. In case of PCR failure, reaction preparation and amplification would have been repeated. After visualisation, PCR replicates were combined and samples belonging to the same collection date were pooled into sub-libraries using different volumes based on strength of PCR products on gels (no visible band = 20 µL, very faint or faint band = 15 µL, visible band =10 µL, bright band = 5 µL) (Alberdi *et al*. 2018). For each sub-library, 1 μL of the PCR positive controls and 10 μL of blanks/PCR negative controls were used. Sub-libraries were cleaned using a double-size selection magnetic bead protocol (Bronner *et al*., 2013) with a ratio of 0.9X and 0.15X of magnetic beads (Mag-Bind^®^ RXNPure Plus, Omega Bio-tek Inc, GA, USA) to sub-library. Two replicates of bead clean-up were performed per sub-library and replicates were individually checked on a 2% agarose gel before pooling.

A second PCR was used to add Illumina tags to each sub-library. Second PCRs were run in duplicate in a final reaction volume of 50 µL using 25 µL of Q5^®^ Hot-Start High-Fidelity 2X Master Mix (New England Biolabs^®^ Inc., MA, USA), 3 µL of each Illumina tag (10 µM; Integrated DNA Technologies, Belgium), 14 µL of molecular grade water (Fisher Scientific UK Ltd, UK), and 5 µL of cleaned sub-library. Second PCR thermal cycling conditions consisted of: initial denaturation at 95°C for 3 min; 8 cycles of 98°C for 20 sec and 72°C for 1 min; final elongation at 72°C for 5 min. Second-round PCR products were checked on a 2% agarose gel alongside their non-tagged, cleaned counterparts to check for size differences after the addition of tags. A second double-size selection bead purification was carried out with a ratio of 0.7X and 0.15X of magnetic beads to PCR products. Tagged, cleaned sub-libraries were quantified using the Qubit™ 3.0 fluorometer and a Qubit™ dsDNA HS Assay Kit (Invitrogen, UK) before being pooled at equimolar concentrations into a single final library. The final library, comprised of 181 eDNA samples and 85 controls, was checked for size and integrity using the Agilent 2200 TapeStation and High Sensitivity D1000 ScreenTape (Agilent Technologies, CA, USA), then quantified with the NEBNext^®^ Library Quant Kit for Illumina^®^ using qPCR on a StepOne Plus real-time PCR platform (New England Biolabs^®^ Inc., MA, USA). Following qPCR, a final dilution to 4nM was performed and 13 pM of the final denaturated library was loaded onto the Illumina MiSeq^®^ with 10% PhiX using 2 × 300 bp v3 chemistry (Illumina Inc., CA, USA).

### Bioinformatics and statistical analyses

Raw sequence reads were demultiplexed using a custom Python script and then processed using metaBEAT (metaBarcoding and Environmental Analysis Tool) v0.97.11 (https://github.com/HullUni-bioinformatics/metaBEAT), a custom bioinformatics pipeline incorporating commonly used open source software. Briefly, Trimmomatic v0.32 (Bolger *et al*., 2014) was used for read quality trimming (phred score Q30). During the trimming step, reads were also cropped to a maximum length of 110 bp and reads shorter than 90 bp were discarded. Additionally, the first 18 bp of remaining reads were trimmed to ensure removal of the locus primers. FLASH v1.2.11 (Magoč and Salzberg, 2011) was then used to merge read pairs into single reads. For subsequent processing, merged reads and high quality forward reads of sequences that failed to merge were kept. A final length filter (106 bp ± 20%) was applied to ensure sequences reflected the expected fragment size (106 bp). Remaining reads were screened for detection of chimeric sequences against our custom reference database for UK fish (Hänfling *et al*., 2016) using the uchime algorithm (Edgar *et al*., 2011), as implemented in vsearch v1.1 (Rognes *et al*., 2016). Clustering at 100% identity in vsearch v1.1 (Rognes *et al*., 2016) was used to remove redundant sequences and possible sequencing errors, and clusters represented by less than three sequences were omitted from downstream processing. Finally, the retained reads were compared against a UK fish reference database (Hänfling *et al*., 2016) using BLAST (Zhang *et al*. 2000) and a lowest common ancestor (LCA) approach based on the top 10% BLAST matches for any query that matched a reference sequence across more than 95% of its length at minimum identity of 100%. Unassigned sequences from this comparison were subjected to a separate BLAST search against the complete NCBI nucleotide (nt) database using the same query and identity parameters.

Final metaBEAT results were summarised as the number of reads assigned to each OTU in each sample screened, henceforth referred to as read counts. The final dataset with the complete fish OTU x read counts matrix was used to run downstream analyses within R v4.1.1 (R Core Team, 2021). A low-frequency noise threshold of 0.001 (0.1%) was applied to the dataset and calculated as the proportion of reads assigned to each fish OTU over the total read counts on a sample-by-sample basis (De Barba *et al*.,2014; Hänfling *et al*., 2016). The choice of the threshold level was guided by the analysis of reads in PCR positive controls that were not assigned to *M. zebra*. For a higher stringency, species-specific thresholds were also applied and the highest number of reads assigned to fish species in PCR negatives and blanks was removed from those species across the entire dataset.

Maps with circles proportional to Arctic charr read counts were used to visualise temporal and spatial patterns at the sites monitored. Maps were created using shape files downloaded from EDINA Digimap^®^ Ordinance Survey service (http://edina.ac.uk/digimap). Shape files were read into R using the package *rgdal* v1.5-25 (Bivand *et al*., 2019) and the *fortify* function together with the package *ggpolypath* v0.1.0 (Sumner, 2016) were used to build the maps. All graphs were plotted using *ggplot2* v3.3.5 (Wickham, 2016). The read counts assigned to Arctic charr on different eDNA sampling dates (all water samples collected) were compared using the non-parametric Kruskal-Wallis test followed by pairwise comparisons between sampling dates using a Wilcoxon rank sum test, where the minimum level of significance was set at *p* = 0.05.

## Results

### Sequencing outputs and bioinformatics

The raw number of sequences generated was 41,160,110. Across all samples/controls, 73% sequences survived the quality trimming step, of which 98% were successfully merged. Following removal of chimera sequences and clustering, the total number of sequences for the library was 16,357,422. After taxonomic assignment against the 12S UK fish database (Hänfling *et al*., 2016), 5,421,189 sequences matched 21 fish OTUs. After application of the thresholds, 18 OTUs were retained in the final dataset (Fig. S1).

### Variation in Arctic charr eDNA signal

Overall, 155,678 reads were assigned to Arctic charr with significant variation across sampling dates (Kruskal-Wallis rank sum test; *χ*^*2*^ = 26.233, *p*< 0.001).

In the pre-spawning period (one sampling event in October 2017), we did not find any reads assigned to Arctic charr at the spawning sites of RN and NTH nor at the sites at OF (Fig. 2).

During the spawning period (between November and December 2017), reads were assigned to Arctic charr for seven sampling events at both NTH and RN (Fig. 2), showing a marked contrast to the pre-spawning period and the post-spawning period (Fig. 2; Table S2). In addition, there was no significant difference between Arctic charr read counts of sampling dates belonging to the spawning season (Table S2). The number of reads assigned to Arctic charr in the spawning period ranged from 30,729 to 182 at RN (shoreline, putative spawning grounds) and from 15,257 to 413 at NTH shore (known spawning grounds). Arctic charr reads were also detected in the deepest waters along the offshore transect during the spawning period (15,689 to 290 at OF1 to OF5; Fig. 2).

The highest read counts for Arctic charr were found at the putative spawning site RN4 in December 2017 (30,729 reads; Fig. 2) and 23 of 55 samples collected at RN during the spawning period were positive for Arctic charr. The highest occupancy and read counts were found in December (7/8 sites, with the highest read counts 30,729 and 13,943 reads at site RN4 and RN8 respectively; Fig. 2). At the known spawning grounds of NTH shore (NTH1, NTH2, NTH3; Fig. 1), the highest read counts of Arctic charr (15,257 reads) were observed at site NTH2 on 13^th^ November 2017 (Fig. 2). All samples collected in autumn showed positive detection of Arctic charr with the exception of sampling site NTH1 on 17^th^ November (Fig. 2). At the deepest sites along the offshore transect (site OF1 to OF5; Fig. 1), the highest read counts matching Arctic charr were found on 8^th^ November 2017 at site OF3 (15,689 reads; Fig. 2). Across offshore sites, 18 samples out of 25 collected showed positive Arctic charr detection across the spawning period (Fig. 2).

In January (after spawning), Arctic charr was only detected at two sites at RN (RN5 = 578 reads, and RN6 = 483 reads; Fig. 2) and at two sites at OF (OF2 = 910 reads, OF5 = 997 reads; Fig. 2). This read count was significantly lower than in the spawning dates of 13^th^ November and 7^th^ December (Wilcoxon rank sum test: 13^th^ November 2017, *p* = 0.0115; 7^th^ December 2017, *p* = 0.0016; Table S2), but not significantly lower than in the spawning dates of 15^th^ and 17^th^ November (Wilcoxon rank sum test: 15^th^ November 2017, *p* = 0.0854; 17^th^ December 2017, *p* = 0.1562; Table S2). In spring (April 2018), Arctic charr was only detected in one sample from the offshore transect (OF3 = 1838 reads; Fig. 2). This read count was significantly lower than the read counts of any sampling events carried out during the spawning period (Wilcoxon rank sum test: 13^th^ November, *p* = 0.0016; 15^th^ November, *p* = 0.0178; 17^th^ November, *p* = 0.0275; 7^th^ December, *p* = 0.0016; Table S2). Arctic charr were not detected at any sites sampled during the final sampling event in July 2018 (Fig. 2).

### Gill-netting survey

Arctic charr spawning individuals were caught and measured on eight out of twelve gill-netting surveys performed in autumn 2017 (Fig. 3). Twelve spawning Arctic charr were caught including one ripe female, four spent or partially spent females, six running males and one spent male (Fig. 3). Males ranged from 25.3 to 31.5 cm in fork length, whereas females ranged from 22.7 to 31.5 cm. On the 8^th^ and 15^th^ of November 2017, the two netting dates overlapping with the eDNA samplings at NTH, one ripe female (30.6 cm) and one spent female (22.7 cm) were caught respectively (Fig. 3).

**Figure 3.**
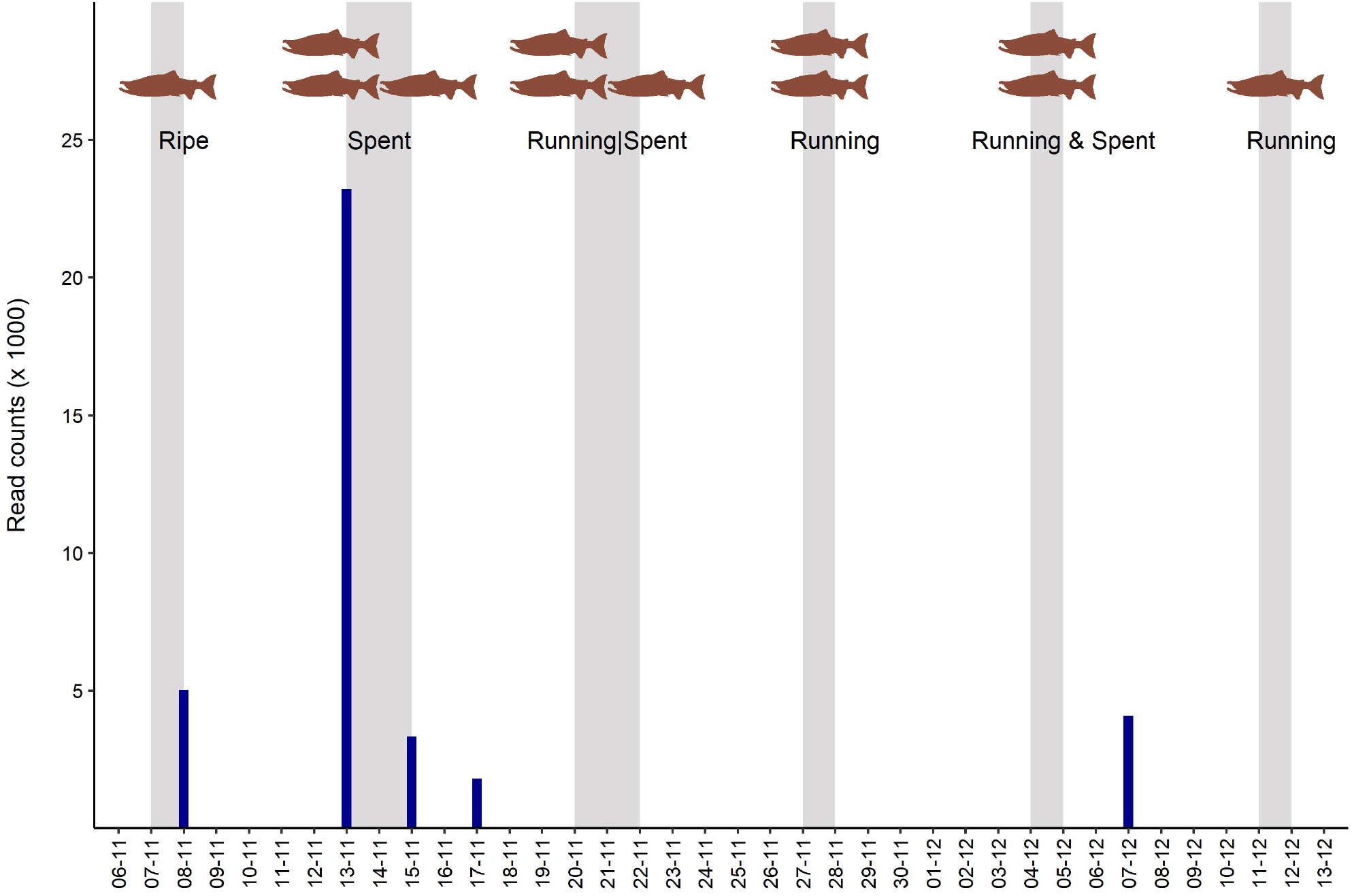
Arctic charr catches and eDNA read counts at the autumn-breeding grounds of NTH. Grey bars indicate the gill-netting dates (net lowered / net lifted) and the number and status of Arctic charr specimens caught. Darkblue bars show the Arctic charr eDNA total read counts from the five water sampling events.

## Discussion

The present study revealed the ability of eDNA metabarcoding to monitor the reproductive activity of Arctic charr autumn-spawning individuals in a lacustrine system. Such molecular observations were supported by catches of mature specimens at the monitored breeding sites where peaks of Arctic charr read counts were detected. In addition, the temporal gradient of the genetic signal observed in autumn 2017 was a further indication of the species’ spawning activity. In this study, we have characterised the times and locations of Arctic charr spawning events, revealing key information on putative breeding localities where spawning has not been monitored or observed for over 50 years.

### Arctic charr eDNA is absent from the monitored sites outside the spawning season

In line with our initial hypotheses, we observed seasonally-limited eDNA detections of Arctic charr at the localities monitored, and Arctic charr eDNA was not found in water samples collected pre-spawning (October 2017; Fig. 2) and post-spawning (January-April-July 2018; Fig. 2) at the shoreline locations of RN and NTH (putative and known breeding grounds respectively). These results confirm previous observations from Windermere showing that Arctic charr is only detected in deep waters outside the species’ spawning season (Hänfling *et al*., 2016; Lawson Handley *et al*., 2019). Such localised distribution of the organisms’ genetic signal is common in lentic systems where the spatial and temporal distribution of eDNA reflects the sites occupied by a species in the water at a given time (Brys *et al*., 2021; Li *et al* 2019; Zhang *et al*., 2020).

Arctic charr feeding grounds are located in the offshore areas of Windermere (Frost, 1977; Mills, 1990); however, the water collection sites along the depth transect (OF; Fig. 1) sampled outside the breeding season showed no detection of Arctic charr (October, April and July; Fig. 2). The limited sampling effort carried out in the deep waters of the lake combined with the aggregated distribution of the species (Jørgensen *et al*., 1993) may have hindered the detection of Arctic charr at the species’ feeding grounds. A more comprehensive sampling effort along the lake midline would likely have found the species in the deepest areas of the lake beyond its spawning season as shown in Lawson Handley *et al*., (2019). In temperate lentic systems, water mixing regimes influence the spatial distribution of eDNA, with eDNA more vertically structured when the water column is stratified (warm seasons) and evenly dispersed when the water column is mixed (cold seasons) (Hänfling *et al*., 2016; Lawson Handley *et al*., 2019). In our previous work we demonstrated that Arctic charr is only detected in mid-depth and lake bottom samples during the summer months (Lawson Handley *et al*., 2019). It is likely that water stratification also contributed to the lower detection of Arctic charr in the deep offshore sites, at least in October 2017 and in July 2018, although possibly also in April 2018 when the thermocline started to develop (ST and EM pers. comm.).

### Arctic charr eDNA is detected consistently at the monitored sites in late autumn

During the Arctic charr spawning season, between November and December 2017, Arctic charr eDNA was consistently detected at offshore (OF) and shallow water sites (RN and NTH; Fig. 2), but exceptional peaks in signal strength were only recorded at shallow water sites (Fig. 2).

The high number of Arctic charr reads found at the shallow breeding habitats of RN and NTH, especially on the 13^th^ November at NTH and 7^th^ December at RN (Fig. 2), exceeded read counts reported otherwise by orders of magnitude. Here, we found an increase in Arctic charr read counts of about 15 and 30-fold compared to other locations and dates within the spawning season (Fig.2). Similar peaks of eDNA were observed during the spawning activity of Japanese eel (*Anguilla japonica*, Temminck and Schlegel, 1846) in a mesocosm experiment when eDNA concentrations were 10 to 100 times higher after the release of gametes (Takeuchi *et al*., 2019). Additionally, Tsuji and Shibata (2021),used a manipulative field experiment, and found that the release of sperm is the main factor explaining peaks in eDNA concentration during fish spawning events. The authors observed 3 to 25 times higher eDNA concentrations during the spawning activity of medaka species (*Oryzias latipes*, Temminck and Schlegel, 1846; *Oryzias sakaizumii*, Asai, Senou and Hosoya, 2012), suggesting that such species-specific peaks in eDNA can be used to identify localities and timings of true spawning events and distinguish them from those sites where other spawning-associated activities occur (Tsuji and Shibata, 2021). Similarly, in our study, the eDNA peak found at NTH on the 13^th^ November coincided with the capture of spent Arctic charr specimens (Fig. 3), whereas the less intense Arctic charr eDNA signal detected consistently at the shoreline breeding sites (RN-NTH; Fig. 2) in autumn could be associated with several reproductive features of the species (i.e. redd-building females, courting and/or competing males, aggregation of mature individuals).

The detection of Arctic charr eDNA at the offshore sites during the breeding season (Fig. 2) might be explained by migratory mature individuals moving from the offshore feeding grounds to the shallow breeding habitats. Time-limited and localised variation in eDNA concentrations have been used to infer fish movements associated to the spawning season in lotic systems whereby visual surveys, egg collection or telemetry have been used to confirm that spatio-temporal variation in eDNA reflects fish migration to the spawning grounds during the reproductive season (Antognazza *et al*., 2019; Erickson *et al*., 2016; Thalinger *et al*., 2019).

Overall and in agreement with previous conventional surveys (i.e. net catches, visual surveys, hydroacoustic, PAM; Bolgan *et al*. 2018; Miller *et al*., 2015; Winfield *et al*., 2015), our results confirmed the suitability of the NTH breeding grounds to support Arctic charr reproductive activity, accounting for (i) the autumn-limited Arctic charr eDNA detections, (ii) the peak of Arctic charr read counts on the 13^th^ November, and (iii) the catches of spent Arctic charr specimens when the eDNA peak was observed (Figs. 2, 3). In addition, given that the same or even higher Arctic charr eDNA read counts were detected at the putative spawning grounds of RN only during the spawning season (Fig. 2), we infer that spawning activity was occurring at this locality even though mature individuals have not been caught at these sites in the last 50 years. Ethical implications of destructive established methods (i.e. gill-netting) restrict the application of these ‘traditional’ monitoring approaches, especially when the target species are threatened and of conservation concern, such as Arctic charr in Windermere (Winfield *et al*., 2009). Therefore, we have demonstrated the suitability of eDNA metabarcoding as a broadly-applicable, non-invasive molecular tool to infer spawning activity through the temporal and spatial localisation of the Arctic charr genetic signal in a large lake.

### Conclusion

In the present study the temporal water sampling coupled with eDNA metabarcoding analysis characterised the Arctic charr spawning activity in a lacustrine ecosystem. We demonstrated that this approach can be used to accurately describe fish spawning locations and timings, to determine the intensity of the spawning effort, and to identify true spawning locations where gametes are released.

As opposed to targeted molecular techniques, the use of eDNA metabarcoding provides an array of “by-catch” community information that can support a more comprehensive evaluation of the status of fish population and ecosystem dynamics. For example, the decline of Arctic charr in Windermere has been linked to the degradation of the species’ spawning grounds as well as to a number of other factors including the establishment of non-native species such as roach (*Rutilus rutilus*, L. 1758) and common bream (*Abramis brama*, L. 1758) (Winfield *et al*., 2008). Such species, facilitated by the changing environmental conditions and eutrophication, have now become abundant or dominant and they are indeed the most commonly detected fish within the lake (Fig. S1; Hänfling *et al*., 2016; Lawson Handley *et al*., 2019). For such reason, we suggest that the periodic use of eDNA metabarcoding in lacustrine ecosystems can assist the monitoring of fish spawning activity and, simultaneously, evaluate changes in fish community composition. The characterisation of these two aspects is essential for an accurate assessment of native fish populations as well as to predict changes in their conservation status.

This research study entails novel applications of eDNA metabarcoding, enhancing the tool’s capabilities far beyond its descriptive and semi-quantitative capacities. Our observations showed that this non-invasive approach provides reliable information on species’ reproductive events, and therefore, eDNA metabarcoding could contribute to the long-term monitoring of fish populations through the spatio-temporal assessment of fish spawning events, an essential aspect for the conservation and management of threatened populations.

## Notes

### Competing Interest Statement

The authors have declared no competing interest.

## References

Adams C., Fraser D., McCarthy I., Shields S., Waldron S., Alexander G. (2003). Stable isotope analysis reveals ecological segregation in a bimodal size polymorphism in Arctic charr from Loch Tay, Scotland. Journal of Fish Biology, 62(2), pp.474–481. https://doi.org/10.1046/j.1095-8649.2003.00044.x

Adams C.E., Bean C.W., Fraser D., Maitland P.S. (2007). Conservation and management of the Arctic charr: a forward view. Ecology of Freshwater Fish, 16(1), 2–5. https://doi.org/10.1111/j.1600-0633.2006.00180.x

Alberdi A., Aizpurua O., Gilbert M.T.P., Bohmann K. (2018). Scrutinizing key steps for reliable metabarcoding of environmental samples. Methods in Ecology and Evolution, 9(1), 134–147. https://doi.org/10.1111/2041-210X.12849

Antognazza C.M., Britton J.R., Potter C., Franklin E., Hardouin E.A., Gutmann Roberts C., Aprahamian M., Andreou D. (2019). Environmental DNA as a non-invasive sampling tool to detect the spawning distribution of European anadromous shads (Alosa spp.). Aquatic Conservation: Marine and Freshwater Ecosystems, 29(1), 148–152. https://doi.org/10.1002/aqc.3010.

Bean C.W., Mainstone C.P., Hall R.A., Hatton-Ellis T.W., Lee A.S.L., Boon P.J. (2018). Guidelines for the Selection of Biological SSSIs. Part 2: Detailed Guidelines for Habitats and Species Groups. Chapter 19 Freshwater Fish. JNCC, Peterborough. https://data.jncc.gov.uk/data/18f27480-a7e1-498c-9cca-f9eacbed2324/SSSI-Guidelines-19-Freshwaterfish-2018.pdf

Bivand R., Keitt T., Rowlingson B. (2019). rgdal: Bindings for the ‘Geospatial’ Data Abstraction Library. R package version 1.4-8. https://CRAN.R-project.org/package=rgdal

Bolgan M., O’Brien J., Chorazyczewska E., Winfield I.J., McCullough P., Gammell M. (2018). The soundscape of Arctic Charr spawning grounds in lotic and lentic environments: can passive acoustic monitoring be used to detect spawning activities?. Bioacoustics, 27(1), 57–85. https://doi.org/10.1080/09524622.2017.1286262

Bolger A.M., Lohse M., Usadel B. (2014). Trimmomatic: a flexible trimmer for Illumina sequence data. Bioinformatics, 30(15), 2114–2120. https://doi.org/10.1093/bioinformatics/btu170

Bracken F.S., Rooney S.M., Kelly-Quinn M., King J.J., Carlsson J. (2019). Identifying spawning sites and other critical habitat in lotic systems using eDNA “snapshots”: A case study using the sea lamprey Petromyzon marinus L. Ecology and Evolution, 9(1), 553–567. https://doi.org/10.1002/ece3.4777

Bronner I.F., Quail M.A., Turner D.J., Swerdlow H. (2013). Improved protocols for Illumina sequencing. Current Protocols in Human Genetics, 79(1), 18.2.1-18.2.42. https://doi.org/10.1002/0471142905.hg1802s79

Brys R., Haegeman A., Halfmaerten D., Neyrinck S., Staelens A., Auwerx J., Ruttink T. (2021). Monitoring of spatiotemporal occupancy patterns of fish and amphibian species in a lentic aquatic system using environmental DNA. Molecular ecology, 30(13), 3097–3110. https://doi.org/10.1111/mec.15742

Corrigan L.J., Lucas M.C., Winfield I.J., Hoelzel A.R., (2011). Environmental factors associated with genetic and phenotypic divergence among sympatric populations of Arctic charr (Salvelinus alpinus). Journal of Evolutionary Biology, 24(9), 1906–1917. https://doi.org/10.1111/j.1420-9101.2011.02327.x

De Barba M., Miquel C., Boyer F., Mercier C., Rioux D., Coissac E., Taberlet P. (2014). DNA metabarcoding multiplexing and validation of data accuracy for diet assessment: application to omnivorous diet. Molecular Ecology Resources, 14(2), 306-323. DOI: 10.1111/1755-0998.12188

Díaz S., Settele J., Brondízio E., Ngo H.T., Guèze M., Agard J., Arneth A., Balvanera P., Brauman K., Butchart S., Chan K., Garibaldi L., Ichii K., Liu J., Subramanian S.M., Midgley G., Miloslavich P., Molnár Z., Obura D., Pfaff A., Polasky S., Purvis A., Razzaque J., Reyers B., Chowdhury R.R., Shin Y-J., Visseren-Hamakers I., Willis K., Zayas C. (2019). Summary for policymakers of the global assessment report on biodiversity and ecosystem services of the Intergovernmental Science-Policy Platform on Biodiversity and Ecosystem Services. https://ipbes.net/sites/default/files/inline/files/ipbes_global_assessment_report_summary_for_policymakers.pdf

Di Muri C., Lawson Handley L., Bean C.W., Li J., Peirson G., Sellers G.S., Walsh K., Watson H.V., Winfield I.J., Hänfling B. (2020). Read counts from environmental DNA (eDNA) metabarcoding reflect fish abundance and biomass in drained ponds. Metabarcoding and Metagenomics, 4, e56959. https://doi.org/10.3897/mbmg.4.56959

Edgar R.C., Haas B.J., Clemente J.C., Quince C., Knight R. (2011). UCHIME improves sensitivity and speed of chimera detection. Bioinformatics, 27(16), 2194–2200. https://doi.org/10.1093/bioinformatics/btr381

Erickson R.A., Rees C.B., Coulter A.A., Merkes C.M., McCalla S.G., Touzinsky K.F., Walleser L., Goforth R.R., Amberg J.J. (2016). Detecting the movement and spawning activity of bigheaded carps with environmental DNA. Molecular Ecology Resources, 16(4), 957–965. DOI: 10.1111/1755-0998.12533

Esteve M. (2005). Observations of spawning behaviour in Salmoninae: Salmo, Oncorhynchus and Salvelinus. Reviews in Fish Biology and Fisheries, 15(1-2), 1–21. https://doi.org/10.1007/s11160-005-7434-7

Franssen J., Blais C., Lapointe M., Bérubé F., Bergeron N., Magnan P. (2012). Asphyxiation and entombment mechanisms in fines rich spawning substrates: experimental evidence with brook trout (Salvelinus fontinalis) embryos. Canadian Journal of Fisheries and Aquatic Sciences, 69(3), 587–599. https://doi.org/10.1139/f2011-168

Freyhof J. and Kottelat M. (2008). Salvelinus alpinus. The IUCN Red List of Threatened Species 2008: e.T19877A9102572. https://dx.doi.org/10.2305/IUCN.UK.2008.RLTS.T19877A9102572.en. Downloaded on 18 October 2021

Frost W.E. (1965). Breeding habits of Windermere charr, Salvelinus willughbii (Günther), and their bearing on speciation of these fish. Proceedings of the Royal Society of London. Series B. Biological Sciences, 163(991), 232–284. https://doi.org/10.1098/rspb.1965.0070

Frost W.E. (1977). The food of Charr,*Salvelinus willughbii (Günther), in Windermere. Journal of Fish Biology, 11(6), 531–547. https://doi.org/10.1111/j.1095-8649.1977.tb05710.x

Garduño-Paz M.V., Adams C.E., Verspoor E., Knox D., Harrod C. (2012). Convergent evolutionary processes driven by foraging opportunity in two sympatric morph pairs of Arctic charr with contrasting post-glacial origins. Biological Journal of the Linnean Society, 106(4), 794–806. https://doi.org/10.1111/j.1095-8312.2012.01906.x

Hänfling B., Lawson Handley L., Read D.S., Hahn C., Li J., Nichols P., Blackman R.C., Oliver A., Winfield I.J. (2016). Environmental DNA metabarcoding of lake fish communities reflects long-term data from established survey methods. Molecular Ecology, 25(13), 3101–3119. https://doi.org/10.1111/mec.13660

Hansen M.J., Krueger C.C., Muir A.M., Klemetsen A., Power M. (2019). Assessing the impact of charr research past, present, and future. Hydrobiologia, 840(1), 1–10. https://doi.org/10.1007/s10750-019-04012-3

Johnson L. (1980). The Arctic charr, Salvelinus alpinus. In: Balon, E.K. (ed.) Charrs, salmonid fishes of the genus Salvelinus. The Hague: Dr. W. Junk Publishers. pp. 15–88.

Jonsson B. and Jonsson N. (2001). Polymorphism and speciation in Arctic charr. Journal of Fish Biology, 58(3), 605–638. https://doi.org/10.1111/j.1095-8649.2001.tb00518.x

Jørgensen E.H., Christiansen J.S., Jobling M. (1993). Effects of stocking density on food intake, growth performance and oxygen consumption in Arctic charr (Salvelinus alpinus). Aquaculture, 110(2), 191–204.https://doi.org/10.1016/0044-8486(93)90272-Z

Kelly R.P., Port J.A., Yamahara K.M., Crowder L.B. (2014). Using environmental DNA to census marine fishes in a large mesocosm. PloS One, 9(1), p.e86175. https://doi.org/10.1371/journal.pone.0086175

Kelly S., Moore T.N., de Eyto E., Dillane M., Goulon C., Guillard J., Lasne E., McGinnity P., Poole R., Winfield I.J., Woolway R.I., Jennings E. (2020). Warming winters threaten peripheral Arctic charr populations of Europe. Climatic Change, 163(1), 599–618. https://doi.org/10.1007/s10584-020-02887-z

Klemetsen A., Amundsen P.A., Dempson J.B., Jonsson B., Jonsson N., O’connell M.F., Mortensen E. (2003). Atlantic salmon Salmo salar L., brown trout Salmo trutta L. and Arctic charr Salvelinus alpinus (L.): a review of aspects of their life histories. Ecology of freshwater fish, 12(1), pp.1–59. https://doi.org/10.1034/j.1600-0633.2003.00010.x

Klemetsen A., Knudsen R., Primicerio R., Amundsen P. A. (2006). Divergent, genetically based feeding behaviour of two sympatric Arctic charr, Salvelinus alpinus (L.), morphs. Ecology of Freshwater Fish, 15(3), pp.350–355. https://doi.org/10.1111/j.1600-0633.2006.00128.x

Klemetsen A. (2010). The charr problem revisited: exceptional phenotypic plasticity promotes ecological speciation in postglacial lakes. Freshwater Reviews, 3(1), 49–74. https://doi.org/10.1608/FRJ-3.1.3

Lawson Handley L., Read D.S., Winfield I.J., Kimbell H., Johnson H., Li J., Hahn C., Blackman R., Wilcox R., Donnelly R., Szitenberg A. (2019). Temporal and spatial variation in distribution of fish environmental DNA in England’s largest lake. Environmental DNA, 1(1), 26–39. https://doi.org/10.1002/edn3.5

Levasseur M., Bergeron N. E., Lapointe M. F., Bérubé F. (2006). Effects of silt and very fine sand dynamics in Atlantic salmon (Salmo salar) redds on embryo hatching success. Canadian Journal of Fisheries and Aquatic Sciences, 63(7),1450–1459. https://doi.org/10.1139/f06-050

Li J., Hatton-Ellis T.W., Lawson Handley L.J., Kimbell H.S., Benucci M., Peirson G., Hänfling B. (2019). Ground-truthing of a fish-based environmental DNA metabarcoding method for assessing the quality of lakes. Journal of Applied Ecology, 56(5), 1232–1244. https://doi.org/10.1111/1365-2664.13352

Low J.J., Igoe F., Davenport J., Harrison S.S.C. (2011). Littoral spawning habitats of three southern Arctic charr (Salvelinus alpinus L.) populations. Ecology of Freshwater Fish, 20(4), 537–547. https://doi.10.1111/j.1600-0633.2011.00502.x

Lynch A.J., Cooke S.J., Deines A.M., Bower S.D., Bunnell D.B., Cowx I.G., Nguyen V.M., Nohner J., Phouthavong K., Riley B., Rogers M.W. (2016). The social, economic, and environmental importance of inland fish and fisheries. Environmental Reviews, 24(2), 115–121. https://doi.org/10.1139/er-2015-0064

Magoč T. and Salzberg S.L. (2011). FLASH: fast length adjustment of short reads to improve genome assemblies. Bioinformatics, 27(21), 2957–2963. https://doi.org/10.1093/bioinformatics/btr507

Maitland P.S., Winfield I.J., McCarthy I.D., Igoe F. (2007). The status of Arctic charr Salvelinus alpinus in Britain and Ireland. Ecology of Freshwater Fish, 16(1), 6–19. https://doi.org/10.1111/j.1600-0633.2006.00167.x

Miller H., Winfield I.J., Fletcher J.M., James J.B., Van Rijn J., Bull J.M., Cotterill C.J. (2015). Distribution, characteristics and condition of Arctic charr (Salvelinus alpinus) spawning grounds in a differentially eutrophicated twin-basin lake. Ecology of Freshwater Fish, 24(1), 32–43. https://doi.org/10.1111/eff.12122

Mills C.A., Heaney S.I., Butterwick C., Corry J.E., Elliott J.M. (1990). Lake enrichment and the status of Windermere charr, Salvelinus alpinus (L.). Journal of Fish Biology, 37(sA), 167–174. https://doi.org/10.1111/j.1095-8649.1990.tb05032.x

Partington J.D. and Mills C.A. (1988). An electrophoretic and biometric study of Arctic charr, Salvelinm alpinus (L.), from ten British lakes. Journal of Fish Biology, 33, 791–814. https://doi.org/10.1111/j.1095-8649.1988.tb05524.x

Piccolo J.J. (2017). The Land Ethic and conservation of native salmonids. Ecology of Freshwater Fish, 26(1), 160–164. https://doi: 10.1111/eff.12263

R Core Team (2021). R: A language and environment for statistical computing. R Foundation for Statistical Computing, Vienna, Austria. URL https://www.R-project.org/

Reist J. D., Power M., Dempson J. B. (2013). Arctic charr (Salvelinus alpinus): a case study of the importance of understanding biodiversity and taxonomic issues in northern fishes. Biodiversity, 14(1), 45–56. https://doi.org/10.1080/14888386.2012.725338

Riaz T., Shehzad W., Viari A., Pompanon F., Taberlet P., Coissac E. (2011). ecoPrimers: inference of new DNA barcode markers from whole genome sequence analysis. Nucleic Acids Research, 39(21), pp.e145–e145. https://doi.org/10.1093/nar/gkr732

Rognes T., Flouri T., Nichols B., Quince C., Mahé F. (2016). VSEARCH: a versatile open source tool for metagenomics. PeerJ, 4, p.e2584. https://doi.org/10.7717/peerj.2584

Sellers G.S., Di Muri C., Gómez A., Hänfling B. (2018). Mu-DNA: a modular universal DNA extraction method adaptable for a wide range of sample types. Metabarcoding and Metagenomics, 2, p.e24556. https://doi.org/10.3897/mbmg.2.24556

Skúlason S., Parsons K., Svanbäck R., Räsänen K., Ferguson M., Adams C., Amundsen P-A., Bartels P., Bean C.W., Boughman J., Englund G., Gudbrandsson J., Hooker O., Hudson A., Kahilainen K., Knudsen R., Kristjánsson B., Leblanc C., Jónsson Z., Öhlund G., Smith C., Snorrason S. (2019). A way forward with eco-evo-devo: an extended theory of resource polymorphism with postglacial fishes as model systems. Biological Reviews, 94, 1786–1808. https://doi.org/10.1111/brv.12534

Smalås A., Amundsen P. A., Knudsen R. (2013). Contrasting life history strategies of sympatric Arctic charr morphs, Salvelinus alpinus. Journal of Ichthyology, 53(10), 856–866. https://doi.org/10.1134/S0032945213100111

Sternecker K., Denic M., Geist J. (2014). Timing matters: species-specific interactions between spawning time, substrate quality, and recruitment success in three salmonid species. Ecology and Evolution, 4(13), 2749–2758. https://doi.org/10.1002/ece3.1128

Sumner M.D. (2016). ggpolypath: Polygons with Holes for the Grammar of Graphics. R package version 0.1.0. https://CRAN.R-project.org/package=ggpolypath

Takeuchi A., Iijima T., Kakuzen W., Watanabe S., Yamada Y., Okamura A., Horie N., Mikawa N., Miller M.J., Kojima T., Tsukamoto K. (2019). Release of eDNA by different life history stages and during spawning activities of laboratory-reared Japanese eels for interpretation of oceanic survey data. Scientific Reports, 9(1), 1–9. https://doi.org/10.1038/s41598-019-42641-9

Telnes T. and Saegrov H. (2004). Reproductive strategies in two sympatric morphotypes of Arctic charr in Kalandsvatnet, west Norway. Journal of fish biology, 65(2), 574–579. https://doi.org/10.1111/j.0022-1112.2004.00432.x

Thalinger B., Wolf E., Traugott M., Wanzenböck J. (2019). Monitoring spawning migrations of potamodromous fish species via eDNA. Scientific Reports, 9(1), 1–11. https://doi.org/10.1038/s41598-019-51398-0

Tsuji S. and Shibata N. (2021). Identifying spawning events in fish by observing a spike in environmental DNA concentration after spawning. Environmental DNA, 3(1), 190–199. https://doi.org/10.1002/edn3.153

Wickham H. (2016). ggplot2: Elegant Graphics for Data Analysis. Springer-Verlag, New York 260pp.

Winfield I.J., Fletcher J.M., James J.B. (2008). The Arctic charr (Salvelinus alpinus) populations of Windermere, UK: population trends associated with eutrophication, climate change and increased abundance of roach (Rutilus rutilus). Environmental Biology of Fishes, 83(1), 25–35. https://doi.org/10.1007/s10641-007-9235-4

Winfield I.J., Fletcher J.M., James J.B., Bean C.W. (2009). Assessment of fish populations in still waters using hydroacoustics and survey gill netting: experiences with Arctic charr (Salvelinus alpinus) in the UK. Fisheries Research, 96(1), 30–38. https://doi.org/10.1016/j.fishres.2008.09.013

Winfield I.J., Hateley J., Fletcher J.M., James J.B., Bean C.W., Clabburn P. (2010). Population trends of Arctic charr (Salvelinus alpinus) in the UK: assessing the evidence for a widespread decline in response to climate change. Hydrobiologia, 650(1), 55–65. https://doi.org/10.1007/s10750-009-0078-1

Winfield I.J., van Rijn J., Valley R.D. (2015). Hydroacoustic quantification and assessment of spawning grounds of a lake salmonid in a eutrophicated water body. Ecological Informatics, 30, 235–240. https://doi.org/10.1016/j.ecoinf.2015.05.009

Winfield I.J., Berry R., Iddon H. (2019). The cultural importance and international recognition of the Arctic charr Salvelinus alpinus populations of Windermere, UK. Hydrobiologia, 840(1), 11–19. https://doi.org/10.1007/s10750-018-3814-6

Zhang Z., Schwartz S., Wagner L., Miller W. (2000). A greedy algorithm for aligning DNA sequences. Journal of Computational Biology, 7(1-2), 203–214. https://doi.org/10.1089/10665270050081478

Zhang S., Lu Q., Wang Y., Wang X., Zhao J., Yao M. (2020). Assessment of fish communities using environmental DNA: effect of spatial sampling design in lentic systems of different sizes. Molecular Ecology Resources, 20(1), 242–255. https://doi.org/10.1111/1755-0998.13105

